# Predicting catchment suitability for biodiversity at national scales

**DOI:** 10.1101/2022.03.31.486513

**Authors:** Barnaby Dobson, Saoirse Barry, Robin Maes-Prior, Ana Mijic, Guy Woodward, William D. Pearse

**Affiliations:** Department of Civil and Environmental Engineering, Faculty of Engineering, Imperial College London; Department of Life Sciences, Faculty of Life Sciences, Imperial College London

**Keywords:** Ecological niche models, Species distribution models, Water quality, Land use, Pollution, invasive species, rare/protected species, biomonitoring, GBIF

## Abstract

Biomonitoring of water quality and catchment management are often disconnected, due to mismatching scales. Great effort and money is spent each year on routine reach-scale surveying across many sites, particularly in the UK, and typically with a focus on pre-defined indicators of organic pollution to compare observed vs expected subsets of common macroinvertebrate indicator species. Threatened species are often ignored due to their rarity as are many invasive species, which are seen as undesirable even though they are increasingly common in freshwaters, especially in urban ecosystems. However, these taxa are monitored separately for reasons related to biodiversity concerns rather than for gauging water quality. Repurposing such monitoring data could therefore provide important new biomonitoring tools that can help catchment managers to directly link the water quality that they aim to control with the biodiversity that they are trying to protect. Here we used the England Non-Native and Rare/Protected species records that track these two groups of species as a proof-of-concept for linking catchment scale management of freshwater ecosystems and biodiversity to a range of potential drivers across England. We used national land use (Centre for Ecology and Hydrology land cover map) and water quality indicator (Environment Agency water quality data archive) datasets to predict the presence or absence of 48 focal threatened or invasive species of concern routinely sampled by the English Environment Agency at catchment scale, with a median accuracy of 0.81 area under the receiver operating characteristic curve. A variety of water quality indicators and land-use types were useful in predictions, highlighting that future biomonitoring schemes could use such complementary measures to capture a wider spectrum of drivers and responses. In particular, the percentage of a catchment covered by freshwater was the single most important metric, reinforcing the need for space/habitat to support biodiversity. We show how our method could inform new catchment management approaches, by highlighting how key relationships can be identified and how to understand, visualise and prioritise catchments that are most suitable for restorations or water quality interventions. The scale of this work, in terms of number of species, drivers and locations, represents a step towards a new approach to catchment management that enables managers to link drivers they can control (water quality and land use) to the biota they are trying to protect (biodiversity).

## 1 Introduction

Water quality is the major determinant of biodiversity in freshwaters globally and in the UK, national biomonitoring schemes have been used for decades to track the impacts of chemical pollutants (Friberg et al., 2011), particularly acidification (e.g., the UK Acid Waters Monitoring Network), organic pollution (e.g., RIVPACS and RICT schemes) and, more recently but to a lesser extent, toxins (e.g., SPEAR classifications). Many of these schemes rely on reach-scale sampling of a subset of the food web (usually macroinvertebrates) and matching observed versus expected biotic indices, based on reference conditions, to infer water quality, largely because direct chemical measures are too costly or insensitive. However, this methodology has some critical limitations and defining suitable reference conditions, for example, in urban areas is challenging or impossible, because these systems are already heavily modified and often contain large populations of multiple non-native invasive species. Non-native species are particularly problematic in this context because they are often excluded from such bioassessment schemes as indicators per se, so their distributional drivers are often poorly understood, even though they will also be sensitive to - and potential indicators of - water quality. The proportions of invasive species in England in our suite of 48 focal species (see Section 2.3) monitored by the Environment Agency is demonstrated in Figure 1 showing this strong urban signature and highlighting their particular potential as indicators in these poorly understood ecosystems. Another major constraint on current schemes is that they tend to take a piecemeal approach to focus on one particular stressor, whereas in reality most freshwaters are exposed to multiple stressors, where disentangling drivers and responses at scale remains a major challenge, especially in urban areas that are exposed to the greatest range of chemical pollutants (M. C. Jackson et al., 2021).

**Figure 1:**
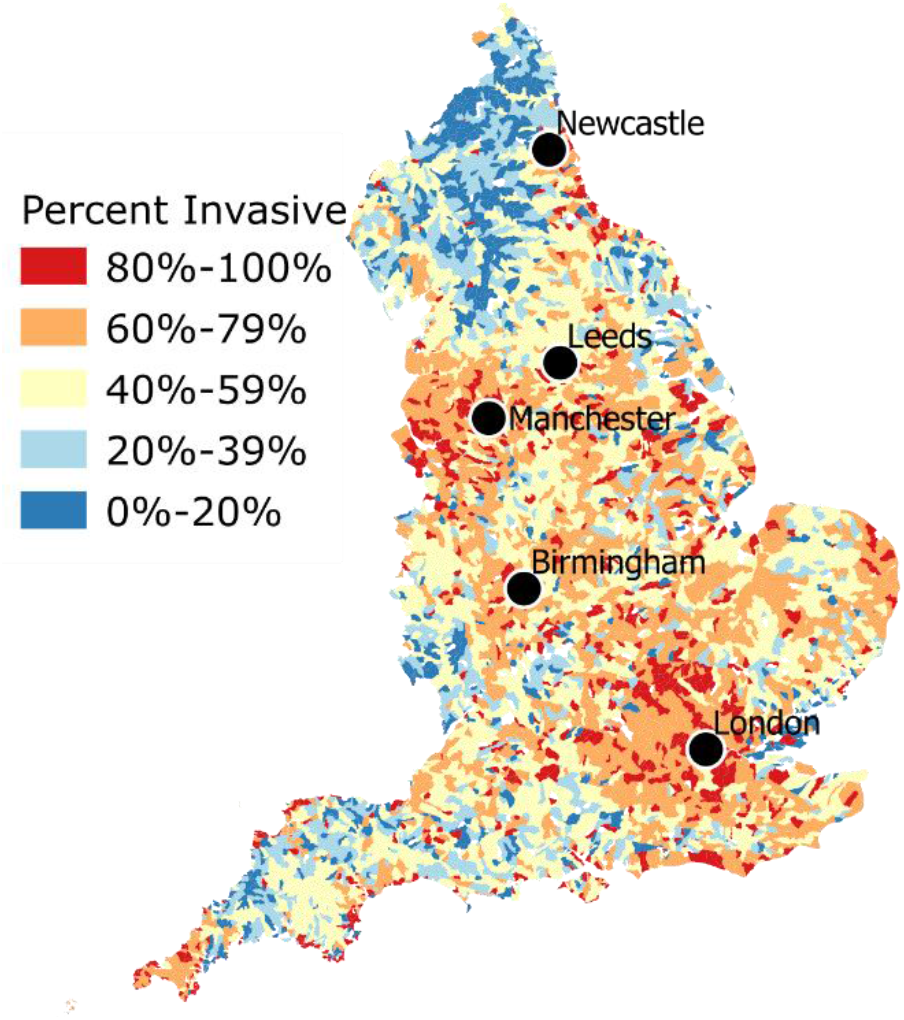
A heatmap depicting the percentage, from our 48 focal species, that are invasive in 5,638 catchments across England. Warmer colours indicates increasing dominance of invasives and selected major conurbations of more than 1.5 million people are highlighted to illustrate the strong urban signal. Species data can be found at Environment Agency (2020a, 2020b), described in Section 2.3. Catchments without samples in either database are omitted.

These limitations and caveats are well-known (Friberg et al., 2011; Harper et al., 2021), but a major bottleneck to addressing them more fully has been the collation and analyses of suitable data that can unpick the drivers of different species’ distributions, especially at larger scales where more meaningful biodiversity management actions can be taken. These limitations also hinder more proactive management, such as restoration, rewilding, and species reintroductions, which are often done in an ad-hoc fashion, with many such efforts failing because the wider context of whether or not the catchment is suitable for the focal species in the first place has not been considered. For instance, if the chemical or physical conditions at catchment scale are prohibitive to the focal species, then restoring habitat patches will yield negligible benefits. Conversely, if managers need to predict the future suitability of particular catchments for invasive or protected species, tools that can map their occurrence across environmental gradients will also be invaluable for prioritising future control and conservation interventions. Traditional biomonitoring schemes are extremely costly and time consuming, in particular because species are so broadly distributed across regions but also responsive to local and fine-scale stressors. Thus, there is a pressing need to determine where and how fine-scaled monitoring data can be matched onto broader catchment scales, and also to gauge the usefulness of citizen science data which are collected in a similar way and are increasingly supplanting routine regulatory surveys.

Current efforts from catchment-scale decision makers towards environmental improvement focus on making in-river conditions central to evaluating the impacts of interventions and for identifying efficient systems-wide decisions (Dobson & Mijic, 2020). However, shaping effective policy for environmental improvements needs to consider biodiversity (Deflem et al., 2021). Without such insights current catchment restorations are likely to remain highly variable and often ineffective (Kail et al., 2015) such that management of pollution in rivers and land use at these scales may be insufficient to protect freshwater biodiversity, and hence vast amounts of time and money may be wasted on unattainable outcomes. To ultimately create meaningful environmental improvements, decision-makers need to be able to target their land-use, infrastructure or agriculture interventions towards predicted changes to biodiversity (Whyte et al., 2020).

Among the tools we can employ to meet this challenge, Ecological Niche Modelling (ENM), sometimes also called Species Distribution Modelling (SDM), aims to link independent variables (presumed ecological drivers) with the presence or absence of species (Guillera-Arroita et al., 2015; Araújo et al., 2019). A range of these studies have demonstrated that water quality samples can indeed be used to predict the presence of specific species of concern (Oliveira et al., 2010; Tamayo & Olden, 2014; Gobeyn & Goethals, 2019; Gallardo et al., 2020; Huml et al., 2020; Kim et al., 2020, 2021). This is a very different approach from the traditional multivariate analyses used in schemes such as RICT in the UK for tracking invertebrate responses to organic pollution, in which an overall index value of ecological status is assigned relative to a “predicted” score based on reference conditions mapped onto that system’s physicochemical attributes and location. An ENM approach could therefore act as a complementary tool, and also a potential validation test, since it can employ independent datasets that are not tied to predefined baselines. Additionally, because these data are often collected for other purposes, it helps maximise the amount of useful ecological information that can be extracted per unit effort or cost.

We propose that the most useful information of this type for decision makers would be at the catchment scale, because this is the scale at which management actions and river basin management plans (RBMPs) are implemented. By using river catchments as the unit for presence-absence sites, rather than more conventional grids or a radius within an observation, Kuemmerlen et al. (2014) demonstrated that predictions benefit from this more informative spatial extent that captures activities across the entire upstream catchment. This tailors information for catchment-scale management and acknowledges that a given river aggregates biotic responses to multiple stressors, and particularly those pollutants that are usually generated on land and at large scales, rather than those solely in the nearfield observation neighbourhood in time and space (e.g., point source toxic spills, sewage outflows). This inherent link with catchment land-use has repeatedly been demonstrated to be one of the strongest predictors of species presence (Dyer et al., 2013; Kuemmerlen et al., 2015), however, to the authors’ knowledge it has not yet been augmented with multiple water quality parameters in species modelling.

Applications across multiple catchments would provide high levels of replication at regional to national scales and, because of the significant differences between catchments, this can reveal strong gradients in multiple stressors that are obscured at smaller scales used in traditional biomonitoring schemes. A national scale approach allows potential drivers and responses to be disentangled in a way not possible using current piecemeal studies. By providing this bigger picture, multi-catchment ENM applications could help to prioritise more targeted follow-up biomonitoring and interventions as and when required, rather than the more ad-hoc (and hence costly) approaches that predominate at present. Another advantage of a catchment-based approach is that the impacts of stressors will be manifested at different scales in space and time, and across different parts of the food web. This allows expanding phylogenetic and functional coverage beyond the more common narrow focus on macroinvertebrates to include multiple trophic levels and even species that span freshwater-terrestrial-marine ecotones (e.g., riparian plants, migratory fishes). Capturing such species that also respond to longer lived and spatially disaggregate stressors can provide more integrated ecological responses than is possible from focusing on a narrow subset of taxa within a small part of the food web.

In this paper, we illustrate how adopting a catchment-scale approach to ENM can provide useful information for decision makers on the relationships between land-use and water quality information with biodiversity conditions. We make this case with four testable hypotheses. First, that water quality and land-use data can be aggregated to meaningful catchment management units and used to make robust predictions on the presence or absence of a given species. Second, that this approach can be scaled to a wide variety of species across the food web, using the same suite of predictors in a simple and succinct framework. Third, that those species can go beyond the traditional biomonitoring indicators to include both those that are under threat as well as those that are non-native invasive species: all of these have niche requirements and so fit the modelling framework we are proposing. Finally, these methods can help managers identify what species, land uses, water quality indicators, and locations should be prioritised for future interventions. In testing the first two hypotheses, we present a method to aggregate species, water quality, and land use information to a catchment-scale and model the presence-absence of species. We demonstrate this tool across all 5,638 Water Framework Directive (WFD) catchments in England as a proof-of-concept for a set of routinely monitored species of concern across the riverine food web, from plants to large predators, and we also present a more detailed case study of a worked example for a focal species, the Atlantic salmon, which is a major concern for nationwide river restoration efforts, with a view to identify what constrains its presence or absence.

## 2 Datasets and pre-processing

A variety of data and pre-processing is required, both in terms of predictors and responses (species observations), to generate presence-absence predictions for different species. All analysis was performed at the catchment scale across England. We outline catchments using the ‘water body’ spatial unit from the WFD. As defined by the European Centre for River Restoration: “Water bodies are the basic unit that are used to assess the quality of the water environment and to set targets for environmental improvements.” Although the UK is no longer an EU member state this framework still bounds its current biomonitoring programmes, and it also serves as a useful test of the potential universality of our approach in terms of being able to ultimately adapt it to the remaining EU countries. In England, the UK region used in this study, there are 5,680 water bodies with an average area of 22km^2^ (124,960km^2^ in total). To avoid confusion, we simply refer to these as ‘catchments’ hereafter. We illustrate the data processing in Figure 2 for example catchments and provide further details in the following sections.

**Figure 2:**
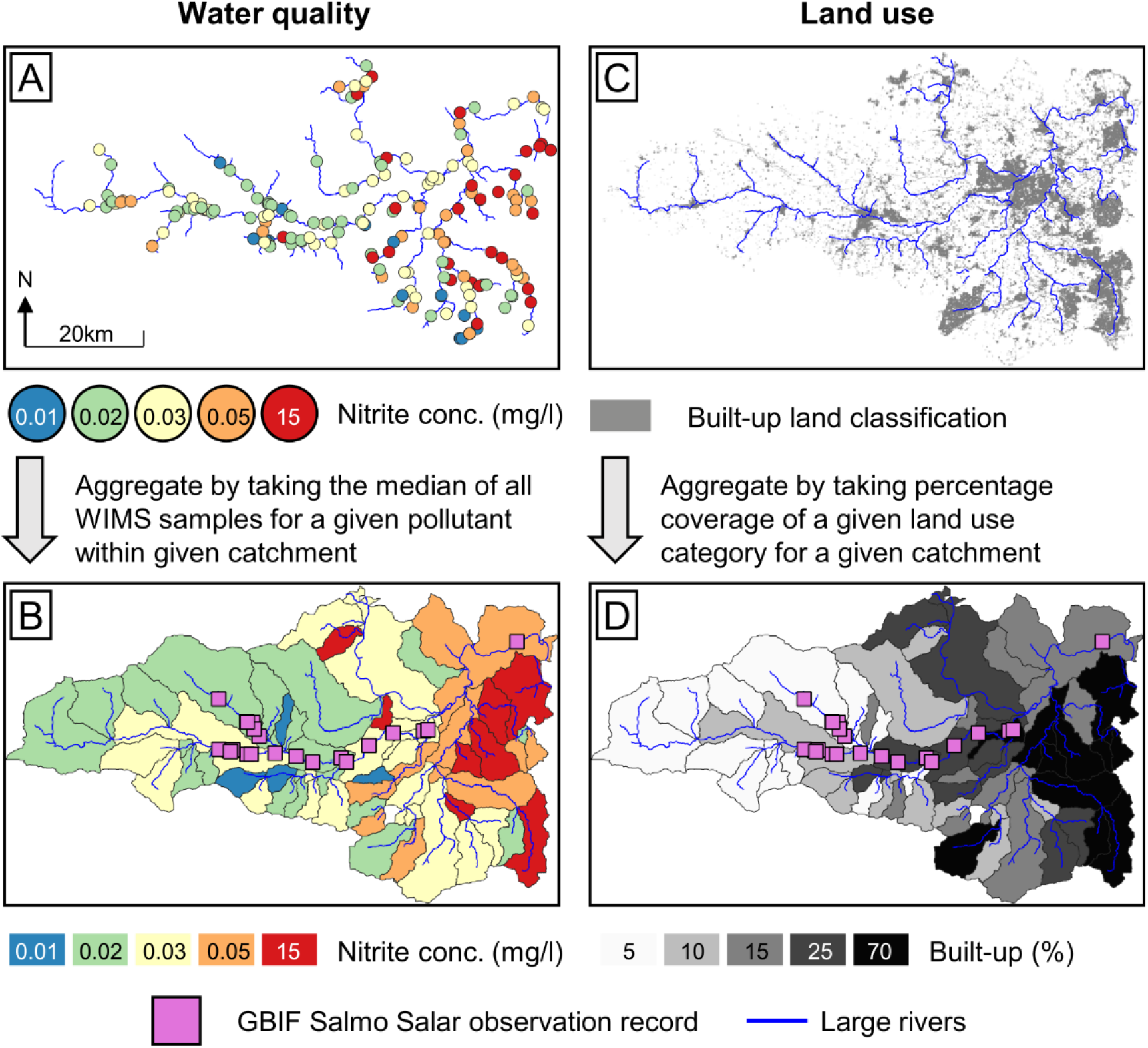
Example of data sources used in this study and their aggregation to catchment scale. The depicted area is of the Kennet, Loddon, and Upper Thames regions in Southern England, these rivers ultimately combine to form the lower urban River Thames. The black catchment outlines in the lower figures are examples of catchments that data is aggregated to in this study (see main text for definition), 58 are depicted here. We note that nitrite concentration and built-up areas are categorised into separate bins in this figure for visual clarity, although the method uses continuous data.

### 2.1 Water quality

The underlying methodology behind catchment management is to assess whether water quality metrics in rivers meet key criteria of use to managers. The English Environment Agency (EA) data that supports this can be found openly available in a single data repository called the ‘Water Data Archive’, commonly referred to as WIMS (Environment Agency, 2020c). Between 2000-2021 (the entire extent of the archive), around 200,000 spot samples of water quality were taken every year and tested for a range of chemical or physical indicators, resulting in an archive containing nearly 60 million individual water quality measurements at 58,000 locations across England. These measurements specify the type of water being sampled (the “sample material”, for example, groundwater, river water or sewage effluent) and thus provide a wealth of information as to the typical water quality profile for water across a range of stages in the water cycle. A breakdown of water quality metric distributions by sample material is provided in Dobson et al. (2021) and has been used to inform recent reports that provide detailed assessment of water quality in both freshwater and estuarine environments (McCormick et al., 2021). The sample material most frequently recorded in WIMS is river water, which, as set out in Section 1, we expect to provide an effective aggregate assessment of overall catchment quality that can be used to predict species presence or absence. Although species often have highly specific niches that they occupy in terms of pollutants (Gobeyn & Goethals, 2019; Deflem et al., 2021), this type of fine-tuned control or monitoring of pollution is not available to catchment managers, who instead rely on a coarse aggregate metric to assess catchment water quality. For example, for a catchment to achieve ‘good’ status under the classification of soluble reactive phosphorus, samples averaged over a year must be less than 50µg P/l (Liu et al., 2022). To reflect this aggregate approach to catchment management, we used the median of all measurements of a given water quality metric within a catchment in the WIMS archive. Figure 2 (A-B) demonstrate the aggregation of nitrite spot samples (2A) over a catchment to a median value (2B) as an example of the general approach.

### 2.2 Land use

Land use data are openly available for the entire United Kingdom (UK) at 25m^2^ resolution from the UK Centre for Ecology and Hydrology (Figure 2C). In this study we use the 2015 land cover map (Owland et al., 2017), considering it the optimal balance between highest resolution and date most representative of the 2000-2019 WIMS data archive period. We note that, aggregated to the average 22km^2^ catchment scale, we expect changes in land use in the 20 years of these data to have minimal impact on our national analysis. Although the data provide up to 21 detailed land class categories defined by (D. L. Jackson, 2000) we use the 10 aggregate classes (broadleaf woodland, coniferous woodland, arable, improved grassland, semi-natural grassland, mountain-heath-bog, saltwater, freshwater, coastal, and built-up areas or gardens), adjudging the differences between aggregate classes to far outweigh the differences within aggregate classes, to maximise the useful information provided to the ENM. To ensure land use information is scale-independent, accounting for different sized catchments, we aggregate by providing the percentage area of a catchment that is accounted for by each land use class (demonstrated in Figure 2D).

### 2.3 Species observations

An openly available source for species observations is the Global Biodiversity Information Facility (GBIF). The API for GBIF can be interacted with automatically via an R package, which we use. GBIF collates both citizen-submitted observations and those from institutions or organisations. The two EA datasets distributed through GBIF are the England Non Native Species Records (Environment Agency, 2020a) and the England Rare and Protected Species Records (Environment Agency, 2020b). Between these two datasets, 232,000 species observations are available for download via the GBIF API. The quality control of these EA data is also listed as ‘high’ because records are provided either directly by biodiversity staff, or through professional surveys. For the purpose of narrowing down a subset of observations to use in this study, we use all species that have been observed in these EA datasets at least 100 times across all sites with sufficient water quality data between 2000-2020 (giving 48 different species and around 220,000 observations). We also test augmenting these EA data with citizen collected data, which doubles the total sample (see Supplement 1).

## 3 Ecological Niche Modelling to reveal ecological drivers at a catchment scale

The overall goal of the ENM in this study was to perform catchment-scale predictions of whether species are present or absent using the independent variables, which will enable variables to be prioritised in terms of how effective they are in making reliable predictions of species occurrences. Ultimately, this will help decision makers improve how they assess the suitability of catchments and whether or not interventions or restoration attempts are likely to be successful.

The workflow demonstrated starts with three raw data sources, the WIMS water quality data archive with a selection of 30 indicators (Section 2.1), a CEH land cover map with 10 categories (Section 2.2), and the 48 focal species downloaded using GBIF (Section 2.3). These data are processed to catchment scale (Supplement S1, see Figure 2 for example) to produce sets of independent variables (water quality and land use) and dependent variables (presence or absence). Catchments are allocated to training or validation sets, and for each species, regression models are fit to training catchments and evaluated on validation catchments (Section 3.1, 4.1, Figures 3-4). We compare independent variable distributions in presence catchments with absence catchments (Section 3.2, 4.2, Figure 5) and map the output of regression models to indicate suitability as a decision-making aid (Section 3.2, 4.2, Figure 6). The presented workflow contains a range of tuneable parameters, described in Supplement 1, and summarised in Table S1. We test different values of these parameters and examine their impact on species predictions performance in Supplement 1.

**Figure 3:**
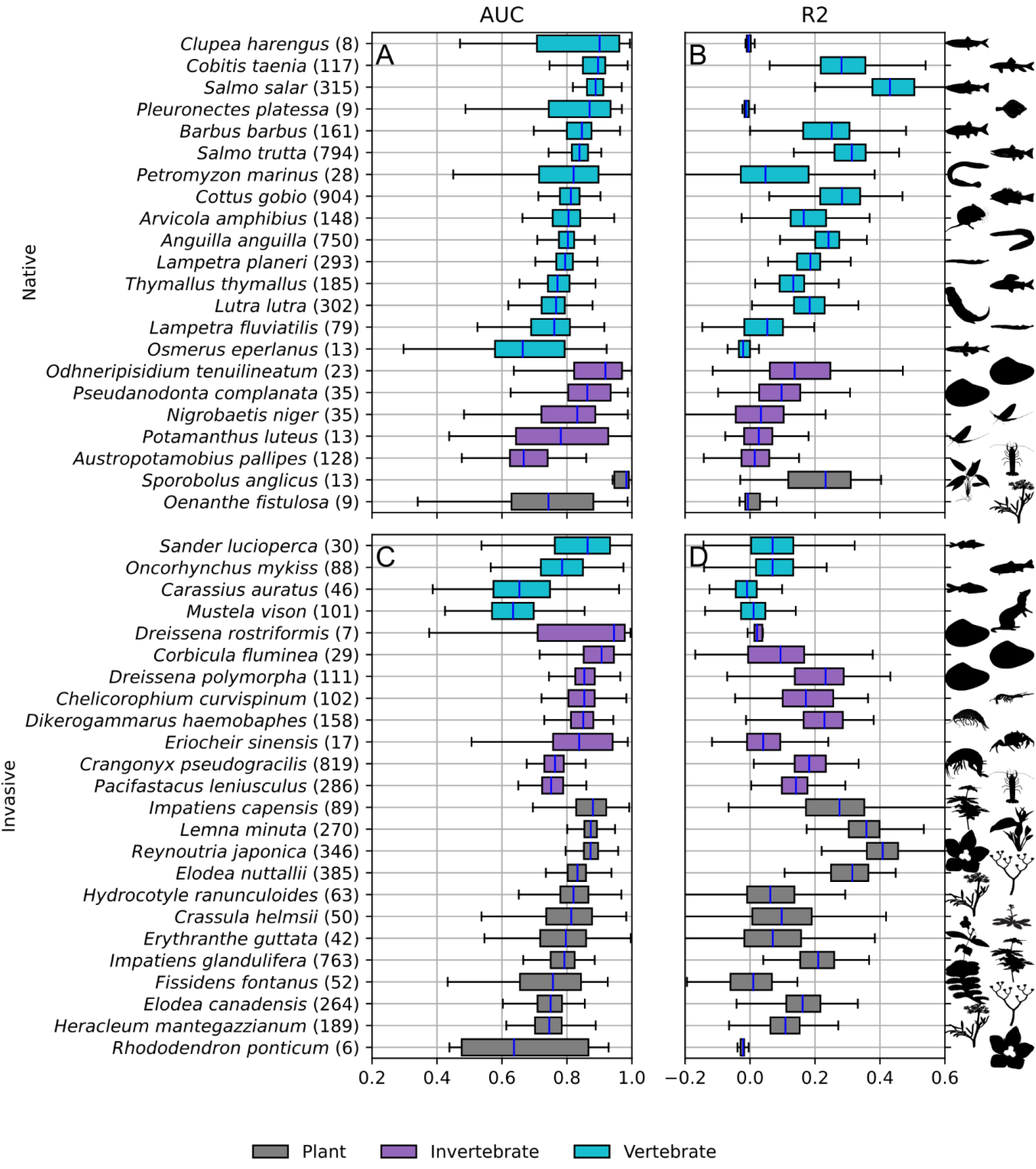
Predictive performance metrics AUC (A, C) and R2 (B, D) for rare and protected native species (A, B) and invasive non-native species (C, D). Boxplot shows range over 100 cross-validation repetitions. Species are grouped by trophic level (indicated by colours). Number of modelled catchments where the species is present is given in brackets next to the species label.

**Figure 4:**
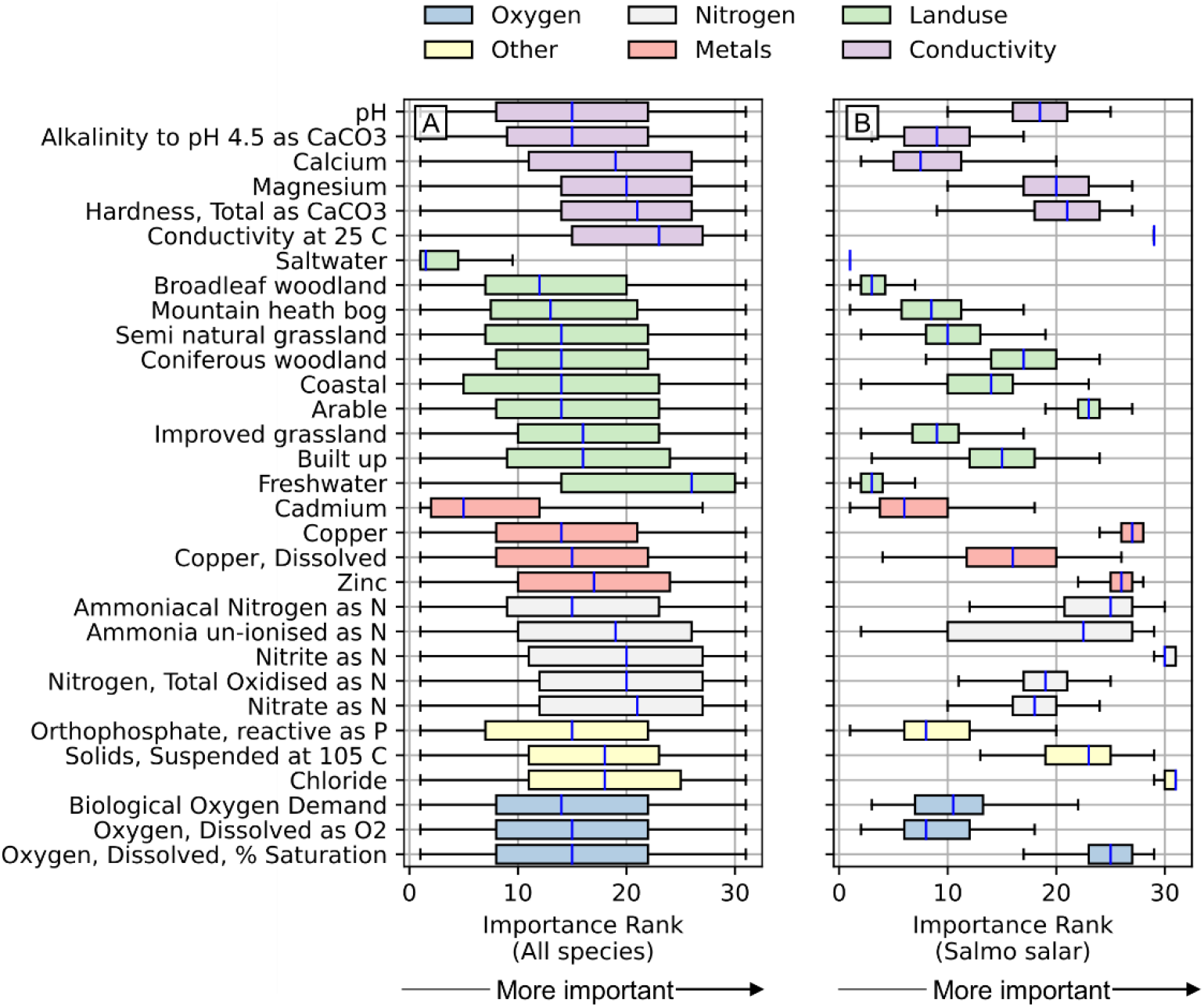
Importance of different independent variables, coloured by broad category. Higher ranks (further to the right on the x-axis) indicate greater importance measured in terms of gain over cross-validation repetitions. (A) Shows results across all species and all cross-validation repetitions. (B) Shows results across all cross-validation repetitions for a single example species (*Salmo salar*).

**Figure 5:**
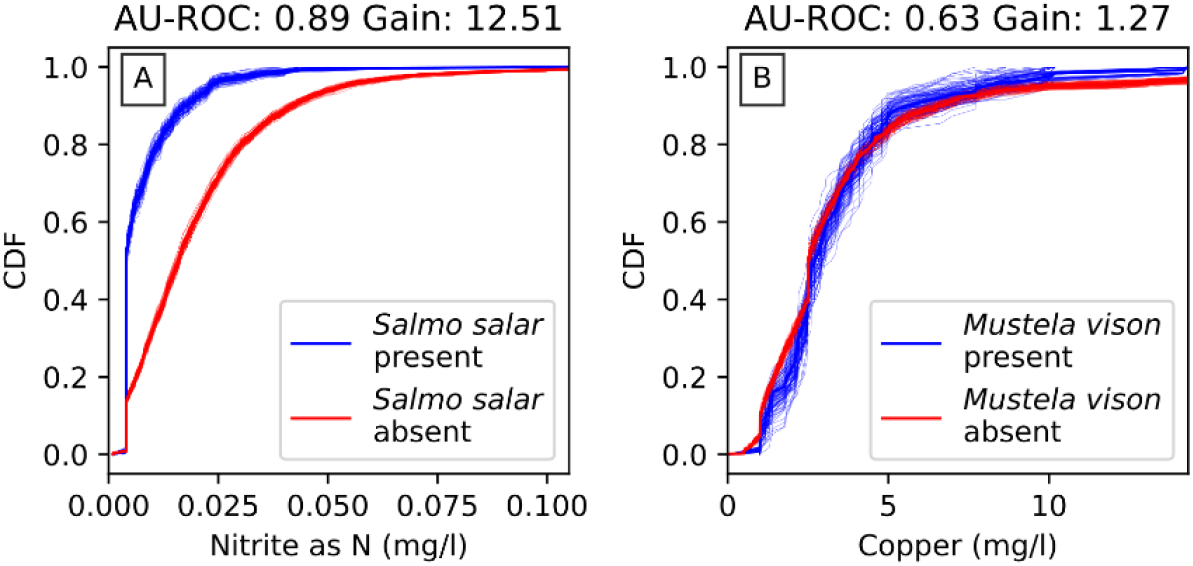
CDFs for water quality metrics for nitrite (A) and copper (B). Red lines show the cross-validation sample distribution in catchments where native Atlantic salmon *Salmo salar* (A) and invasive American mink *Mustela vison* (B) are absent, and blue lines in catchments where both these predator species are present.

**Figure 6.**
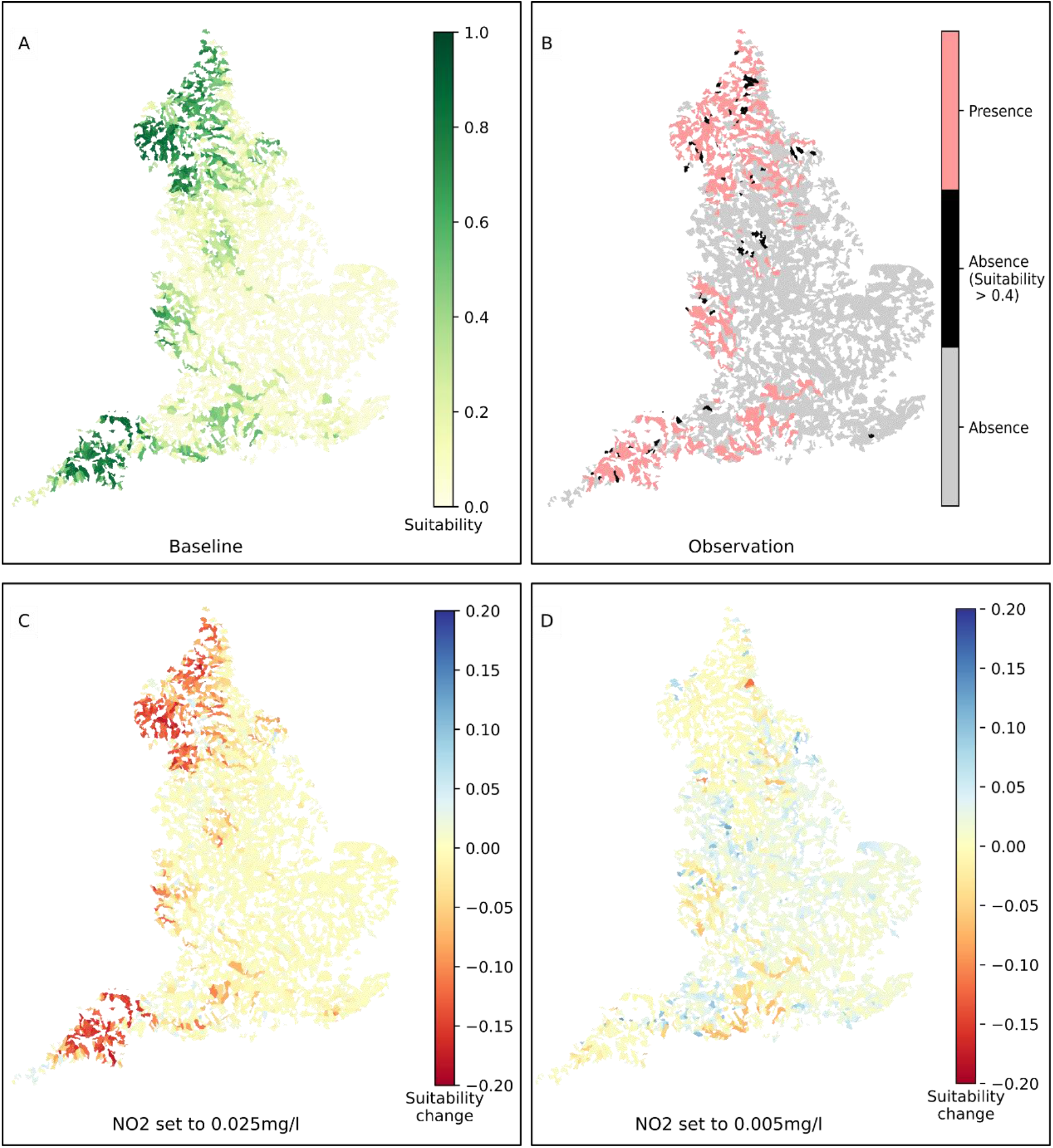
English catchments (water bodies) coloured by (A) the suitability for Atlantic salmon (*Salmo salar*) as predicted by 100 random forest regression models, (B) the presence or absence of salmon with high suitability water bodies that have no salmon observations highlighted, (C) the change in suitability if nitrite concentration (NO2) was set to a high value (0.025mg/l) in every water body, (D) and a low value (0.005mg/l). Water bodies with insufficient water quality indicator samples were omitted.

### 3.1 Regression modelling to predict presence-absence

A range of approaches exist to fit independent variables to dependent variables in the context of ENM; indeed, any statistical method falling under the regression or classification family is viable. In this study, we use a random forests approach because it has had widespread recent application in the water sector (Tyralis et al., 2019) and also in ENM (De Simone et al., 2021). The method is also robust to overfitting, which is a risk in this study because of the high number of independent variables (30 water quality indicators and 10 land use classes). The Python package XGBoost (Chen & Guestrin, 2016) contains support for random forests regression and was used in this study. XGBoost creates regression models fitting with and without including specific independent variables and measuring how much performance is gained with inclusion. This relative performance gain, ‘gain’ hereafter, can be considered a measure for how important an independent variable is in predicting dependent variables. Recording the importance of independent variables is critical in highlighting the relationships that matter for a decision maker.

To evaluate the performance of our proposed method in a rigorous manner, cross-validation was implemented by omitting data from the regression fitting process and evaluating the trained regression models on these omitted data. For each species, catchments were used for either training or validation; the XGBoost random forests regression model was fit to the training catchments and evaluated on the validation catchments, recording the area under receiver operating characteristic curve (AU-ROC), the R^2^ value, and the gain for each independent variable. In an ideal experiment, this process would be repeated using every catchment individually as the validation catchment. As a compromise for computational effort, 10% of the data was randomly allocated into a validation set of catchments with the remainder used for training, and this process was repeated 100 times for each species to bound the likely true performance metrics and independent variable importance (measured by performance gain).

### 3.2 Understanding ecological drivers

The outputs of the random forests regression are predictive performance metrics for each species and gain values for each independent variable for each species. The combination of this information can direct decision makers’ attention towards potential drivers that affect ecological endpoints. For example, a species with high performance metrics can be well explained by the information contained in the independent variables, and those variables with the highest gain will matter the most. These high performance, high gain relationships are those of most potential use to a decision maker. The approach of investigating potential species-independent variable relationships by examining distributional differences between present sites and absent sites has been demonstrated in previous studies (Harrow-Lyle & Kirkwood, 2021). In this study, by plotting the distribution, in terms of a Cumulative Distribution Function (CDF), of the independent variable in sites where the species is absent and comparing against sites where the species is present, the profile of a catchment that is suitable for a given species can be identified. As with the cross-validation of the random forests, we plot the CDF for randomly sampled sites 100 times to capture the variability.

Because our method uses openly available data at a national scale, results can be mapped over all catchments with sufficient water quality data. Because the output of each regression model is a number varying between 0 (absence) and 1 (presence), we can use these models as indicators of suitability for a given species. This enables independent variables to assess species suitability in regions that have not been well surveyed for biodiversity information. A suitability map also provides more granularity and thus greater utility to a decision-maker in comparison to solely a presence/absence classification. The models can also be used to estimate the change in suitability that results from a change in independent variables, thus exploring the potential of different future intervention options.

## 4 Results

### 4.1 Predicting species presence using water quality and land use data

In Figure 3, we present the distribution of performance metrics across cross-validation repetitions by species, grouped by class. All classes contained some species with high performance metrics and some with low. In general, high R2 correlates with a high AU-ROC, but not exclusively. R2 values are sensitive to the total number of present sites (with rarer species being hardest to achieve high R2), while AU-ROC values are not. Thus, we use AU-ROC as the primary performance metric. Figures 3A-B show results for rare and protected native species, while Figures 3C-D show results for invasive and non-native species. There was negligible difference in performance between predictions for native species and invasive (both with an average AU-ROC 0.81), although both categories contained a mix of difficult to predict and easy to predict species. For 90% of modelled species, predictions can be made with ‘fair’ model skill (AU-ROC > 0.7 (Hosmer et al., 2013)) and for 60% with ‘good’ model skill (AU-ROC > 0.8).

The distribution of ranked gain values (i.e., gain in performance metric when that independent variable was included in the regression model) highlights the importance of different independent variables aggregated over all species over all cross-validation repetitions (Figure 4A). Ranks were used because gain values are not comparable between different species. Each category contained important variables and less important variables, although nearly every variable was ranked as either of high or low importance for some species. The independent variables of nitrite concentration, conductivity, hardness, and the percentage of catchment that is the freshwater land category accounted for the highest median importance ranks, and thus as the most important independent variables. In general, the conductivity category measurements (purple) and nitrogen-based pollutants (grey) were the most important.

Although useful for revealing broad trends, grouping all pollutants together masked important differences for some individual focal species. We provide the same information but for a selected species’ (Atlantic salmon, *Salmo salar*) cross-validation repetitions only (Figure 4B), which revealed a different picture from the importance plot for all species (4A). Here the range in an independent variable’s importance was significantly reduced and just a few metrics (nitrite, chloride, conductivity, copper and dissolved oxygen) stood out as consistently important. Because *Salmo salar* was also one of the best predicted species (Figure 3), we can expect that these variables are able to explain its presence/absence and thus that these relationships are significant.

### 4.2 Identifying relevant information for decision makers

The method enables decision makers to investigate potential species-independent variable relationships in more detail. In Figure 5A we plot the CDF of nitrite in catchments where *Salmo salar* is present and the CDF in catchments where it is absent, we select this relationship because Figure 3 highlights that *Salmo salar* are well predicted and Figure 4 shows that nitrite is high gain independent variable for *Salmo salar* models, thus we would expect the relationship to be especially important. Figure 5A shows that catchments with *Salmo salar* have significantly lower nitrite levels than those without, regardless of the cross-validation repetition. In contrast, to demonstrate an unimportant relationship, Figure 5B plots the CDF of copper for catchments with the invasive top predator, American mink, *Mustela vison* (which Figure 3 demonstrates is one of the worst predicted species) either present or absent, showing no significant differences between the two CDFs. A check of the scientific literature can help to elucidate potential mechanisms behind these results and in the *Salmo salar* case, for example, nitrite is well known to be toxic to salmonids, supporting Figure 5A, and it is ameliorated by chloride (see importance of chloride Figure 4) (Eddy et al., 1983). In contrast the mink, *Mustela vison* is insensitive to comparable levels of copper in its native range (Aulerich et al., 1982), suggesting that it may be operating under different constraints as an established invasive species with no natural enemies in the UK.

Figure 6A shows the output of fitted regression models, i.e., a suitability indicator, for *Salmo salar* for catchments that have sufficient water quality indicator samples, we contrast this with Figure 6B, which shows solely presence or absence. By highlighting catchments with high suitability but no *Salmo salar* observations (black catchments, Figure 6B) we also show the most promising candidates for catchment restoration.

In Figures 6C-D we modelled a scenario where we set nitrite to a high (0.025mg/l) and low (0.005mg/l) concentration for every catchment, re-evaluated suitability, and plotted the resulting change in suitability. This type of approach can be employed by planners to identify how changes to pollutant concentrations could alter suitability of a given species in the landscape. In Figure 6C increasing nitrite to a concentration that is prohibitive to *Salmo salar* (Figure 5A shows that only 5% of catchments with *Salmo salar* present have a median nitrite concentration higher than 0.025mg/l) will dramatically reduce the suitability of many catchments across the UK (many deep red catchments). In contrast, Figure 6D shows that reducing nitrite to a lower concentration (Figure 5A shows that around 50% of catchments with *Salmo salar* present have median nitrite concentration around 0.005mg/l) does improve suitability (most catchments are light blue), however it is not sufficient alone to dramatically change the suitability landscape for *Salmo salar*. Considering the substantial cost of implementation, this reduction in permitted nitrite concentration could be ruled out as a useful legislative option for conservation of *Salmo salar*.

## 5 Discussion

This study reveals the huge potential for capitalising on and repurposing different forms of biomonitoring data to assess biodiversity and the potential for catchment management at regional to national scales for a wide range of species spanning the entire food web. It also shows how such data can guide management of both native species and exotic invaders, and even for using the latter as viable indicators of water quality, especially in degraded urban ecosystems where invaders often predominate.

The first hypothesis presented in this work was that water quality and land-use data can be aggregated to catchment scale for making predictions on species’ presence-absence. The high performance metrics observed in Figure 3, whose robustness was confirmed through the extensive cross-validation, give strong support for this hypothesis. From a catchment management perspective, these results are highly encouraging, as water quality and land use are often tractable variables that can be shaped by interventions at individual catchments. By equipping policymakers with the tools to understand what species-suitable catchments look like, we could tailor water quality regulations in a more targeted and efficient manner and open dialogues between communities, policy, and the water industry to help make decisions based on a significantly firmer evidence base.

The second hypothesis was that this approach could be applied using the same suite of water quality and land use variables for all species. A key finding of this study is that nearly all variables provide important information at some level, but the relative importance of each variable differs across species (Figure 4). That means there is a large amount of complementary insights that can be gleaned by going beyond simpler, single-species based or macroinvertebrate-assemblage perspectives to one that considers many species from across both the Tree of Life and the food web. From a management perspective this means that it is desirable to continue monitoring a wide variety of variables to maximise the sensitivity of the monitoring scheme. This will be challenging considering the dramatic reduction in sampling effort that has taken place over the last 20 years in England alone (a more than 50% decrease in pollutant records in the WIMS data archive when comparing 2000-2005 to 2015-2020 (Environment Agency, 2020c)). However, it is encouraging that citizen science data (easily available through the GBIF API that was used), which increasingly makes up for the growing shortfall in routine sampling, can also be applied in our approach. Further, having a high percentage of freshwater land cover within a catchment being the most important variable for predictions (Figure 4A) suggests that giving wildlife space, above all, must be a key priority for planners. This supports the conclusions of Lawton (2010) in the influential “Making space for Nature” report that is now the cornerstone of the UK’s 25 Year Plan for the Environment. Our results show that, although Lawton focussed mostly on terrestrial ecosystems, the same principles also clearly apply in the freshwater realm, providing a clear path to link data, models, and policy.

Answering the third hypothesis showed that it is possible to make use of even seemingly “undesirable” non-native species to provide us with additional indicators of water quality. ENM is often challenging if a species is so widespread that their tolerances and limits are unclear (they are everywhere), whereas if a species is rare then there are few occurrences from which to produce a model (see (Guillera-Arroita et al., 2015; Araújo et al., 2019) for a broad review of ENM use and fitting). Invasive species, like native species, fall along this spectrum, but they also present the additional challenge of often being excluded from, or marked negatively against, assessments of ecological quality. This is problematic in areas where ecosystems still provide functions and services but invasives are common and reference conditions are lacking: notably, lowland urban ecosystems. Given that most of the world’s human population lives in urban environments, and most cities are on river floodplains, with the UK being no exception, an ability to include invasive species in assessment of ecosystems is a major benefit. Thus, we were surprised and heartened to find that we were able to produce accurate models of non-native species’ distributions, and that we could find no correlation between predictive accuracy and overall occupancy. This suggests that current monitoring approaches, which are mostly driven to focus on indicators of more “pristine” or observed/reference conditions, could be quickly and easily augmented with additional metrics based on non-natives. As such, our approach could serve dual purposes – it can use these species as indicators in their own right and also provide information to managers about their potential spread based on their environmental tolerances and niche requirements. In addition, the effectiveness of the ENM suggests that it could be used to augment ongoing long-term monitoring as well as *priori* identification of watersheds that are prone to invasive species. This enables biomonitoring to adopt a far more predictive and hence proactive approach than has been the case to date.

The final hypothesis was that these methods can help managers identify what species, land uses, water quality indicators, and locations should be prioritised for future interventions. In Figures 5A-B we demonstrate how the proposed methodology reveals which independent variables have significantly different distributions in catchments where a species is present than catchments where a species is absent. These relationships can then be verified by literature study to target catchment management more effectively. The fitted regression models for a given species can be used to scale up decision making by creating suitability maps (Figure 6A) to identify the best potential restoration sites (Figure 6B) hence to project the estimated impacts of interventions (Figure 6C-D). We ultimately envisage pairing this method into water decision making so that managers can directly target the physical and chemical water indicators that will have the greatest biodiversity benefits, rather than relying on proxy criteria for water quality.

A variety of parametric choices and statistical methods were tested in Supplement 1, Figure S1. This highlighted the role that Citizen Science (CS) observations may play in ENM, more than doubling our observation sample size although, unlike with the Environment Agency data, we recommend bias correction with CS data (Isaac et al., 2014; Isaac & Pocock, 2015) to maintain predictive performance. There are a variety of other statistical refinements that could be made to improve the proposed approach still further, including pre-processing of independent variables to account for collinearities (for example, principal components analysis) and to remove redundant information (De Marco & Nóbrega, 2018). Further investigation to sensitivities could also help account for species that are sensitive to the extremes in catchment water quality indicators rather than the median across samples. Additionally, more sophisticated treatment of pseudo-absence information has also been shown to improve predictions based on citizen-collected data (Iturbide et al., 2015; Domisch et al., 2016). We suggest that the proposed method can accommodate these refinements and our analysis presented in Supplement 1 provides a simple framework to investigate their impacts.

Overall, we argue that approaches such as the one presented in this paper can be used to make efficient and effective use of data that is commonly gathered for other purposes, to give an overview of the bigger picture of constraints on the distributions of key species – both native and exotic – at larger scales where major policy and management interventions are often conducted. It could also be used to prioritise more targeted biomonitoring at reach scales using traditional techniques and offers promise for integration with emerging technologies and tools, such as eDNA (Carraro et al., 2018), which are especially well-suited to presence/absence occupancy data at scale. This could augment existing schemes as well as provide a means for developing new tools for monitoring, modelling, and managing our freshwaters to mitigate the impacts of the multiple emerging stressors they will face in the coming decades.

## 6 Conclusion

In this paper, we used land use data and water quality indicator samples to predict the presence-absence of species. We used regression modelling to make predictions at a catchment scale, adjudging that this scale is the most relevant to decision makers. Our results revealed that it is possible to make accurate and robust predictions of a variety of species, both invasive and rare native, using common and open access data in catchments across England. The proposed method can be used to create tailored decision-making information, as we demonstrate in an example with Atlantic salmon. This information includes identifying water quality indicators of interest and understanding their distribution in catchments with Atlantic salmon, as well as using mapping techniques to identify suitable catchments where they are missing and estimating changes in suitability that could be engineered through interventions. Ultimately, we hope that this method will enable catchment managers to better target interventions towards improving freshwater biodiversity.

## Acknowledgements

The research reported in this paper was taken as part of the CAMELLIA project (Community Water Management for a Liveable London), funded by the Natural Environment Research Council (NERC) under grant NE/S003495/1. The authors are grateful to Collette Taylor for comments on the manuscript that have improved the paper.

