## Supplement 1 for "Predicting catchment suitability for biodiversity at national scales"

**Summary**

In this supplement we describe the evaluations performed to explore the impacts of parametric choices on performance of ENM.

### Supplement 1

#### S1 Overview

We tested our presented method from the main paper using different values for 12 tuning parameters, Table S1. Using a one-at-a-time sampling regime, we tested each value shown in Table S1 with other parameters set to their default values (bold) for 100 cross-validation repetitions, as in the main paper. This both allowed us to investigate some useful scientific questions, e.g., is there a role for Citizen Science data and does it need bias correction, or to examine how sensitive the method is to choice of regression modelling parameters.

| Parameter | Summary | Values |
|---|---|---|
| use.public | Species observation data sources | True (Citizen collected + EA), False (**EA only**) |
| list.length | Remove short observation lists | **0.0**, 0.5, 1.0, 2.0, 4.0 |
| site.visits | Remove poorly sampled catchments | **0.0**, 1.0, 2.0, 5.0, 10.0 |
| quantile | Water quality catchment aggregation | 0.025, 0.25, **0.5**, 0.75, 0.9 |
| N.retained.catchments | Ensure all catchments have water quality samples | Fill, 500, 750, **1000**, 1250 |
| P.sub | Percentage of catchments to use in training | 50, 60, 70, 80, **90** |
| XGB.estimators | Number of trees | 10, **40**, 80, 160, 320 |
| XGB.sub | Subsample ratio of training | 0.5, 0.6, **0.7**, 0.8, 0.9 |
| XGB.depth | Maximum tree depth | 3, 4, 5, **6**, 7 |
| XGB.gamma | Minimum loss reduction to partition a tree | 0.0, 0.5, **1.0**, 2.0, 5.0 |
| XGB.col.tree | Subsample ratio of columns for trees | 0.6, 0.7, 0.8, 0.9, **1.0** |
| XGB.col.level | Subsample ratio of columns for levels | 0.4, 0.55, **0.7**, 0.85, 1.0 |

Table S1. A list of all parameters tested in this study and values that they take. Parameters used in the main study are highlighted in bold.

#### S2 Parameters

S2.1 Water Quality

S2.1.1 Aggregation to catchment scale

We tested a variety of percentiles ($5^{th}$, $25^{th}$, $50^{th}$, $75^{th}$, $95^{th}$) to aggregate water quality indicators, because we expect some species to prefer higher values of some water quality indicators (e.g., dissolved oxygen) and some to be prefer lower values for others (e.g., nitrate in otherwise nutrient-limited areas).

S2.1.2 Treatment of missing samples

Having a water quality sample for every catchment is a prerequisite for most regression methods. Thus, we tested an approach where missing data were filled with the median value for that pollutant across all catchments, and an approach where pollutants were removed from analysis until a pre-specified number of sites had a sample for every site for every water quality indicator. We started with the 30 most frequently measured water quality indicators in the WIMS database, however the actual number of indicators used was always less then this to ensure every site had a value. Different parameter values for this number of sites threshold were tested, described in Section 3.

S2.2 Species observations

S2.2.1 Citizen collected data

We also tested citizen-science data because it is growing in popularity and has the potential to be applied at global scales. To do this, for each of these 48 species, we downloaded any other observations of them that exist in the study region, which increased the total number of observations to around 484,000, more than doubling our original dataset. However, because citizen science data may have less rigorous quality control and potentially subject to more biases than EA surveys (Isaac and Pocock 2015; Troia and McManamay 2016), we tested a variety of approaches in the ultimate observation dataset to use including: EA data only, all data, and all data augmented with bias correction techniques as described and defined in (Isaac et al. 2014).

S2.2.2 Bias correction

A range of biases can occur in observational data (Isaac and Pocock 2015). Bias correction techniques are typically used to ensure that presence data points are correct with greater certainty (Isaac et al. 2014). This process is complicated by studies such as this that aim to predict both presences and absences (rather than simply studying presence locations, as is most common); a questionable observation may raise uncertainty on the validity of assuming presence, but equally we could not reasonably infer the inverse (absence) from a questionable observation either. Thus, in experiments where bias correction would suggest absence in a site with a questionable observation, we instead omit the site from that repetition.

We test two forms of bias correction: list length and site visits, both defined in detail in (Isaac and Pocock 2015). For an observation to be included in a study, list length bias correction requires it to be made on sites and days with a total number of observations greater than a given threshold. The aim of this is to remove opportunistic observations, i.e., observations in which an observer has noted a single species and reports only that species. These opportunistic observers are considered less likely to be conducting a rigorous sampling survey and their data are considered questionable. To account for the variable sizes of catchments, we set the threshold for inclusion as a per-22km$^2$ value (selected because it is the average catchment size). For example, an observation must come from a list with 2 or more observations per 22km$^2$ on a given day in a given catchment to be included. In our default experiment, we perform no list length bias correction, but we test a range of values (0.5, 1, 2, 4 observations per 22km$^2$).

Site visit bias correction requires a site to have been visited more than a certain number of times to be included in a study. The aim of site visit bias correction is to remove sites that are poorly sampled (relative to their size) on the basis that observations will provide a less representative picture of the catchment. As with the list length bias correction, we set a per-22km$^2$ threshold for total number of visits that a site must have received for its observations to be included in an experiment. Again, in our default experiment, we perform no site visit bias correction, but we test a range of values (1, 2, 5, 10 visits per 22km$^2$).

### S3 Results

We present the median R2, and median AU-ROC averaged across all species over all cross-validation iterations for different tuning parameter samples in Figures S1 and S2.

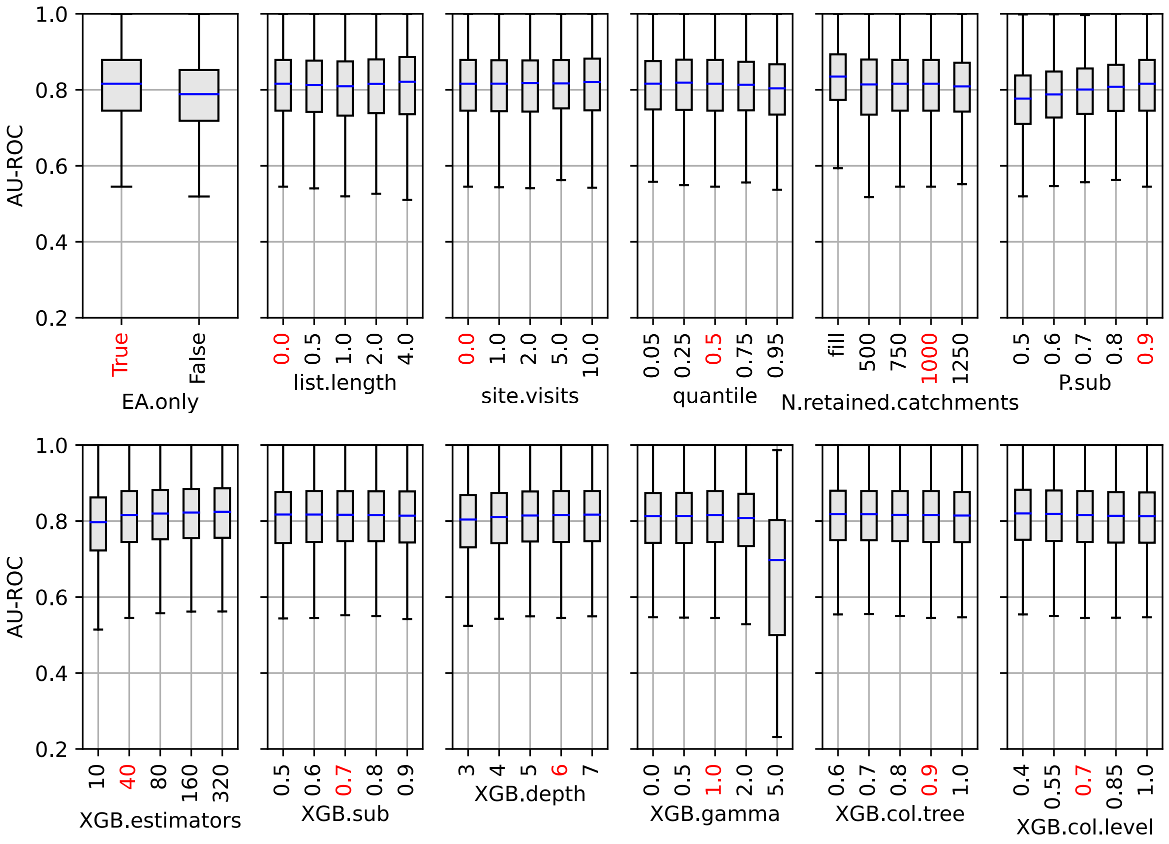


Figure S1: The distribution of performance metrics (AU-ROC) across cross-validation repetitions for different tuneable parameters. The value shown in red is the value used in the main study.

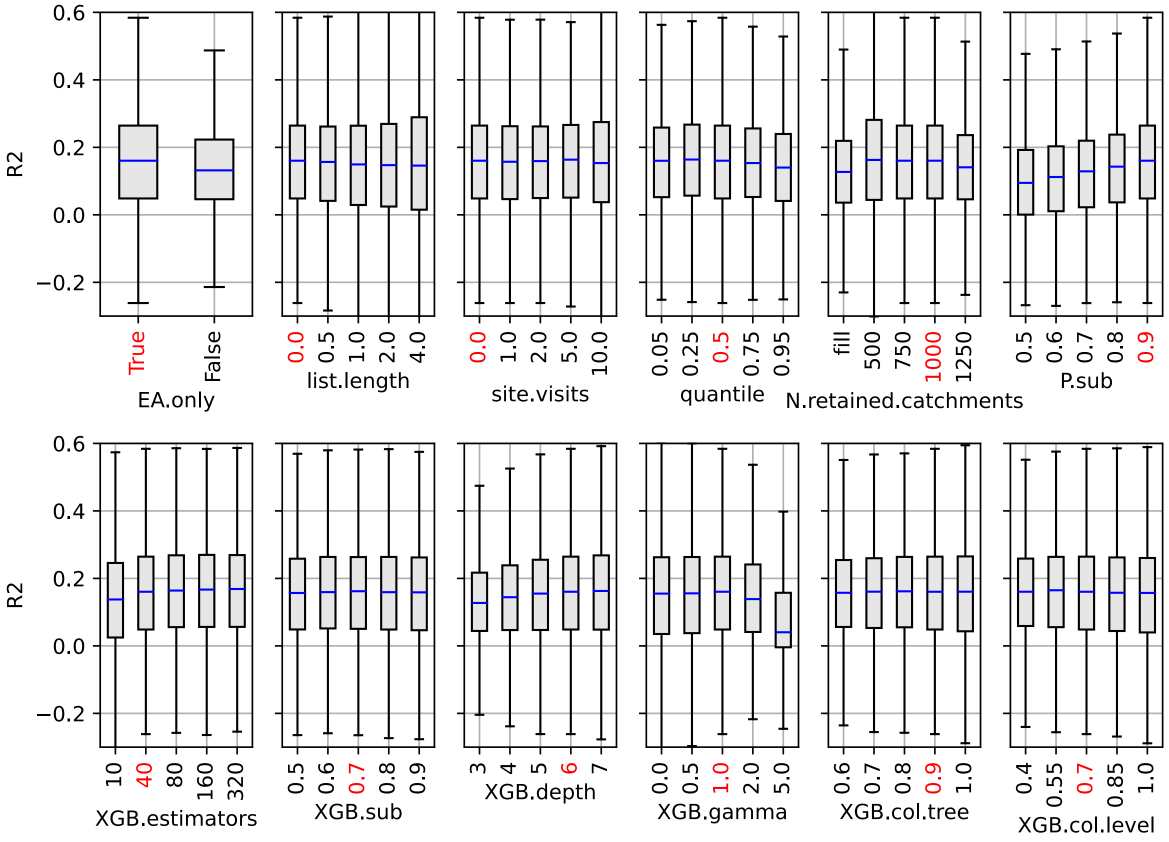


Figure S2: Same as S1 but showing R2.

Using the EA observations only (AU-ROC 0.81, R2 0.16), rather than the entire citizen-science supplemented observation dataset (AU-ROC 0.79, R2 0.13) proved increased performance metrics. This effect dissipated when the entire dataset was used in conjunction with bias correction (AU-ROC 0.81, R2 0.15). Bias correction made no significant difference when applied only to the EA observations. We note that, although these differences are small, they are averaged across 100 cross-validation repetitions and all 48 species.

Model performance was sensitive to the choice of quantile to which the catchment scale water quality data was aggregated. Specifically, at the 95th percentile performance degraded (AU-ROC 0.80, R2 0.13). Water quality samples exhibit high levels of variability, so these more extreme percentiles are unstable and not representative of typical catchment conditions.

Other tuning parameters had smaller impacts on performance. Parameters that reduced overfitting improved performance, levelling off once overfitting had been prevented. Parameters that reduced the amount of training data (particularly the amount of presence samples) reduced performance and increased variability.
